# Isoform-Level Analysis of 10x Genomics Single-Cell cDNA Libraries from Cultured K562 Cells Using Long-Read Sequencing

**DOI:** 10.1101/2025.08.12.668929

**Authors:** Jörg Bachmann, Remi-André Olsen, Agrima Bhatt, Nick Crang, Orlando Contreras-López, Anja Mezger

**Affiliations:** Department of Biochemistry and Biophysics, Stockholm University, Science for Life Laboratory, Stockholm, Sweden; Division of Gene Technology, KTH Royal Institute of Technology, Science for Life Laboratory, Stockholm, Sweden; Department of Cell and Molecular Biology, Karolinska Institutet, Science for Life Laboratory, Stockholm, Sweden

## Abstract

Integration of Oxford Nanopore Technologies (ONT) long-read sequencing with 10x Genomics single-cell cDNA libraries enables novel transcript detection, isoform analysis and captures full-length gene body coverage. The purpose of the study was the comparison of three approaches for sequencing 10x Genomics Chromium Single Cell cDNA libraries using long-read sequencing: single-cell full-length transcript sequencing by sampling (FLT-seq), the cDNA-PCR Sequencing Kit (SQK-PCS111) and the PCR Expansion Kit (EXP-PCA001). Our aim was to evaluate their efficiency in enriching full-length cDNA fragments, identifying barcodes, detecting novel isoforms and mutations, and characterizing transcript coverage profiles.

## Introduction

Single-cell transcriptomic methods have given insight into cellular identity and function in health and disease. With focus at the level of the cell, differential gene expression in complex tissues can be analysed. 10x Genomics Chromium Single Cell Gene Expression is a widespread method, applicable for a wide range of cell types and encompassing the whole workflow from cell suspension to the analysis of transcriptomic data with cellular resolution.

In Chromium Single Cell Gene Expression, full-length mRNA is first captured from fresh cell or nuclei suspensions, methanol fixed or cryopreserved cells (*Figure 1A*). Then, a cell-specific molecular barcode is added, and a cDNA library is generated via reverse transcription and template switching. For short-read sequencing, these cDNA libraries are enzymatically fragmented in order to fit the requirements for Illumina sequencing. This means that while the method yields single-cell resolved gene expression, the full-length information of the mRNA is lost by fragmentation and with that, the possibility to identify novel mRNA isoforms, differentially expressed isoforms, and mutations/SNVs.

**Figure 1.**
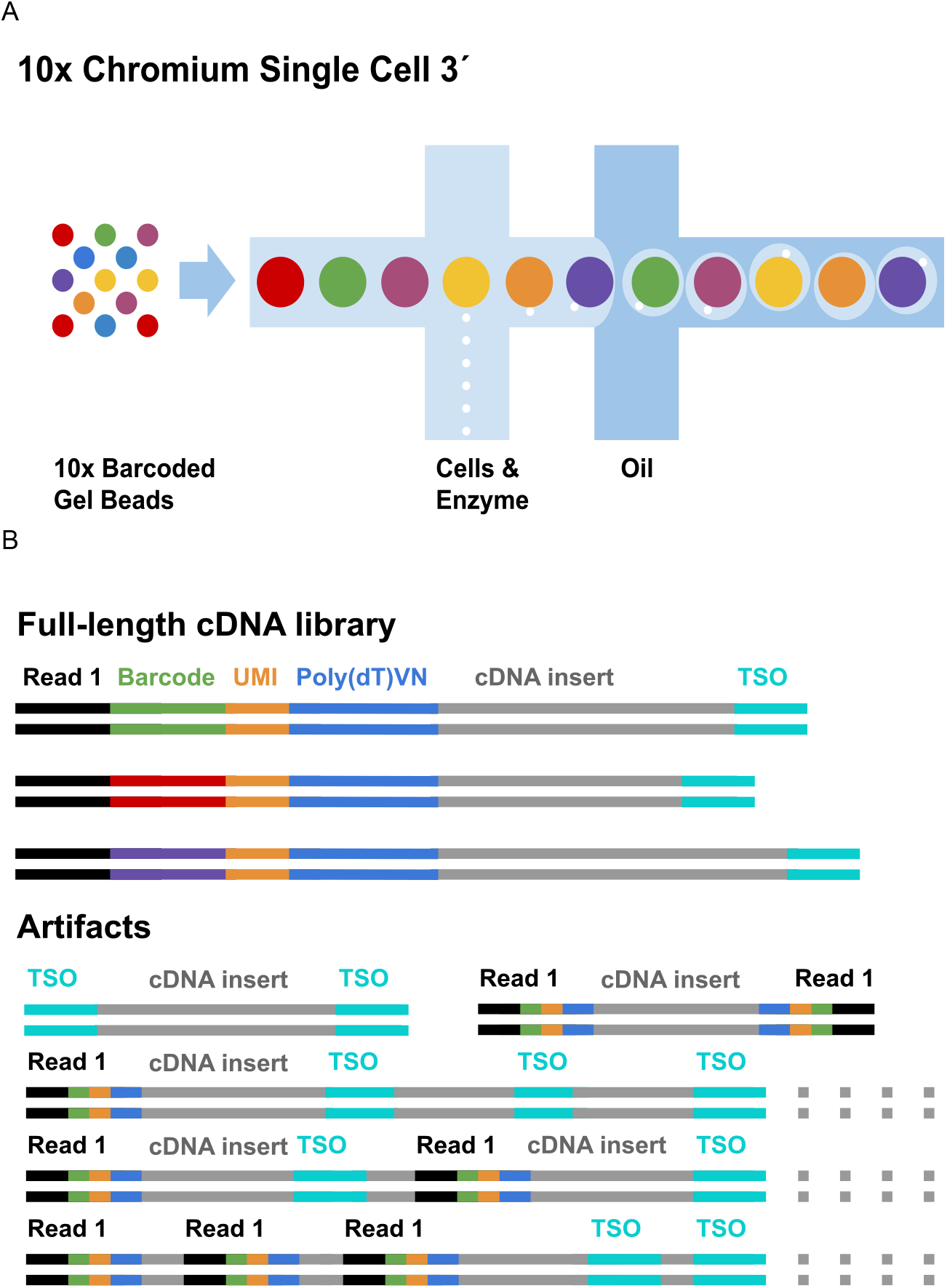
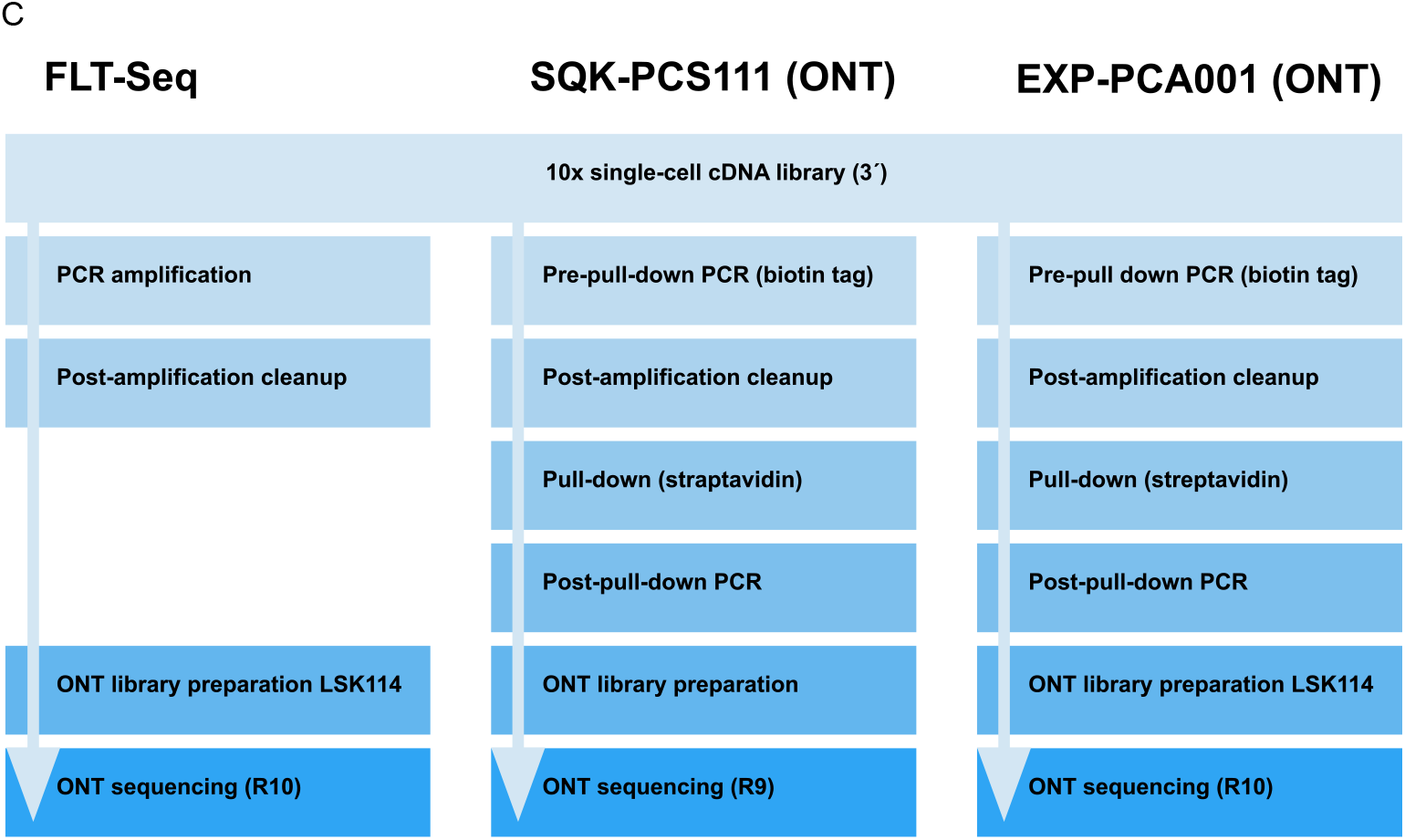
Overview of methods. **A)** cDNA library construction with 10x Chromium Single Cell Gene Expression (adapted from 10x) **B)** Full-length cDNA and different artifacts (adapted from 10x) **C)** Methods for ONT library construction with enrichment of cDNA library

Recently, methods that integrate third-generation long-read sequencing with single-cell transcriptomics have been described to sequence full-length mRNA and yield different splicing isoforms; these “single-cell RNA isoform sequencing” methods are reviewed in Gupta *et al*. 2024.

In the Chromium Single Cell Gene Expression cDNA library, generated through reverse transcription and template switching, a substantial portion of the library can consist of artifacts caused by incorrect template switching, known as template switching oligo (TSO) artifacts (*Figure 1B*). Long-read sequencing technologies typically sequence native DNA without amplification, with sequencing adapters added via ligation. However, applying this approach directly to the cDNA library results in a high prevalence of TSO artifacts in the final dataset. To mitigate this, the cDNA library must first be enriched for the desired fragments, those containing the structure: read 1 - barcode - UMI - poly(dT) - transcript cDNA - TSO, prior to the addition of sequencing adapters.

We evaluated long-read sequencing of 10x Genomics Chromium Single Cell cDNA libraries to assess the feasibility of incorporating mRNA isoform and mutation detection capabilities. We used Oxford Nanopore Technologies (ONT) for the long read sequencing and compared three different methods for sequencing the full-length cDNA libraries: “single-cell **f**ull-**l**ength **t**ranscript **seq**uencing by sampling” **FLT-seq** (Tian *et al* 2021), and two protocols available through ONT: using either the cDNA-PCR sequencing kit **SQK-PCS111** or the PCR expansion kit **EXP-PCA001 (***Figure 1C, Table S1*)

The specific aims of the pilot study were to establish the sequencing depth needed for the barcode identification and in a bigger picture to validate long-read sequencing as an “add-on” to Chromium Single Cell to detect mRNA isoforms and mutations. Publicly available FLT-seq data (“scmixology2”, accession: ERR9958136) was analysed as a reference point for data quality (Tian *et al* 2021). The study was structured around four key research questions:

1. Can full-length cDNA libraries be enriched effectively using these approaches?
2. To what extent can novel isoforms and multi-transcript loci be detected using long-read sequencing?
3. What is the profile of gene body coverage across transcripts, and how does it compare to reference data?
4. What is the isoform length distribution captured by each method?

## Results & Discussion

### Enrichment of full-length cDNA libraries

All three methods in this pilot study aim to enrich full-length cDNA libraries while limiting artifacts (*Figure 1C*). Since the desired cDNA libraries are flanked by the Read 1 sequencing primer and the TSO at each end, we can use the average location of the TSO from the read ends to compare the performance of the methods. In the relative distance of TSO to read ends, our FLT-seq data (in orange) performs very similarly to the publicly available FLT-dataset (in red) (*Figure S1*). Our PCA001 (green) data performs slightly better than the FLT-seq in terms of average distance of TSO from read ends, while the PCS111 data shows a higher average distance of TSO from read ends, as well as a higher variation in relative distance. In absolute distance, TSO is located closer than 200 bp from the read ends in >88% of reads for all methods. (*Figure S2*).

In addition to the variation in average distance of TSO from read ends, we observed reads with multiple TSO annotations in all tested methods, and to a lesser degree, reads with multiple sequences of the Read 1 sequencing primer. We quantified repeated occurrences of TSO and Read1 sequences within the reads across four datasets: PCS111, FLT-seq, PCA001, and scmixology2. The percentages of reads containing two or more TSO hits were 4.0%, 14.6%, 2.3%, and 4.1%, respectively. Similarly, the percentages of reads with repeated Read1 hits were 3.4%, 7.7%, 2.1%, and 3.4%, respectively (Table S2).

### Pipeline results

All three methods in our dataset yield similar results when analyzed with the “wf-single-cell” (v. 1.0.3) workflow from epi2me-labs (*Figure 3*). The workflow can roughly be characterised by four consecutive steps:

1. Reads that fail to meet the QC criteria of one step are not passed on to the subsequent step.
2. 86.1-89.8% of reads are retained after adapter trimming (compared to 85.8% for the public reference FLT-seq dataset).
3. 66.2-74.5% of reads are successfully assigned to valid cell-specific barcodes (compared to 73% for the public reference FLT-seq dataset).
4. In terms of gene expression metrics, our three datasets show slightly lower performance compared to the reference FLT-seq dataset. Specifically, the percentage of reads assigned to genes ranges from 51.2% to 58.7% in our datasets, versus 60.3% in FLT-seq. Similarly, transcript assignment rates are 28.3% to 32.6%, compared to 42.1% in the reference dataset. This discrepancy may reflect biological differences between datasets: the scmixology2 reference contains an equal mixture of cells from five human cancer cell lines (H2228, H838, H1975, HCC827, A549; Tian *et al*., 2021), each with distinct gene expression profiles, whereas our dataset was derived from a single cell type.

Standing out is the method PCS111, which shows a lower barcode matching efficiency than the other methods. (*Figure 2*). This difference likely reflects the fact that the older R9.4.1 nanopore chemistry (97.3% accuracy) used for the PCS111 protocol generally yields lower sequencing accuracy than the more recent R10.4 chemistry (98.3% accuracy) used for the other methods (Ni *et al*., 2023).

**Figure 2.**
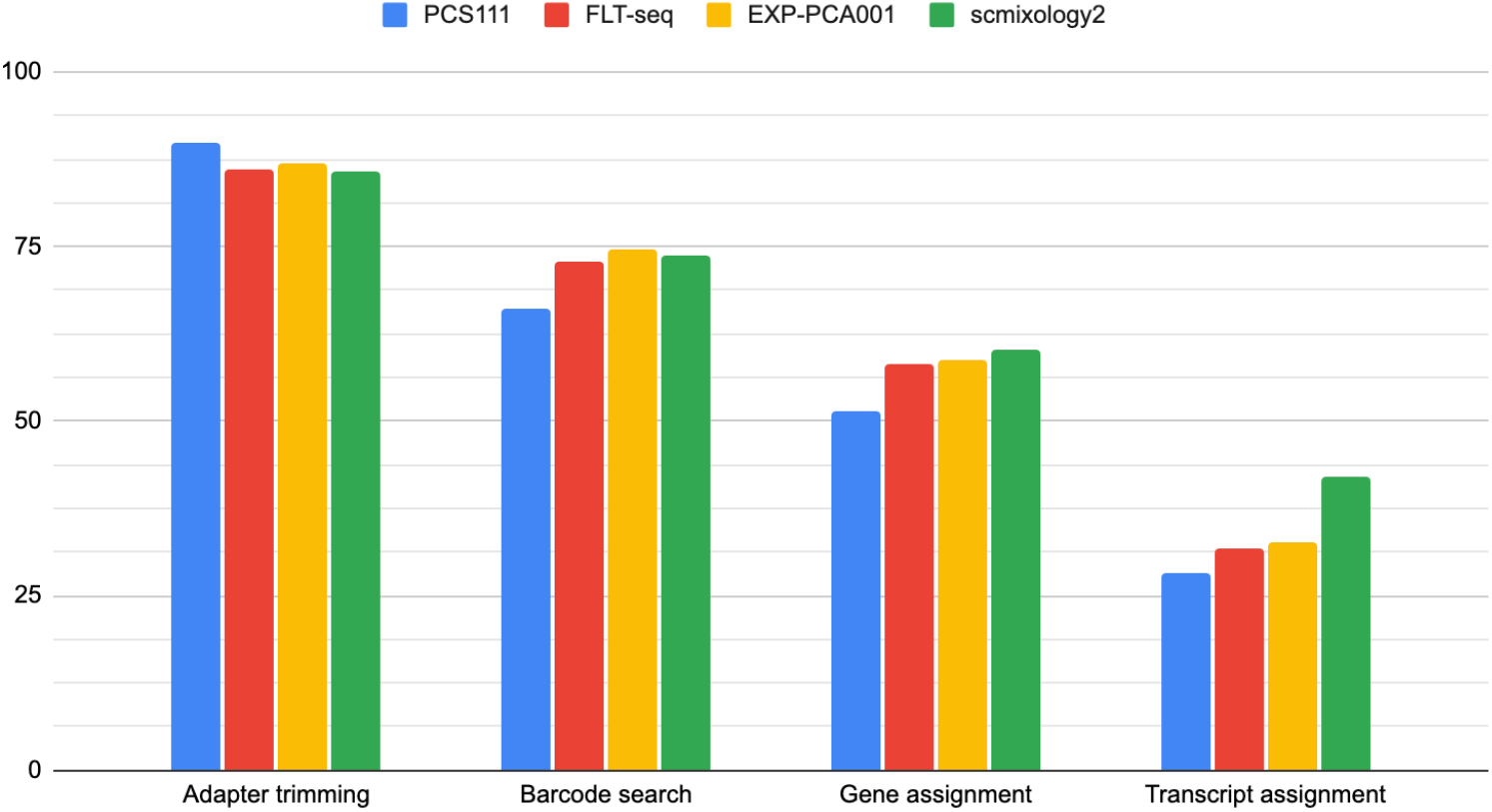
The percentage of reads passing each step of the wf-single-cell pipeline. In succession from left to right, *e.g*. reads matching cell barcodes or reads matching annotated genes.

#### Isoform results

The isoforms expression strongly correlate between the three methods (*Figure 3A*), with a Pearson correlation coefficient of 0.83-0.85 for the pairwise comparison of the normalised transcript levels.

In the subset of exact and complete intron-chain matching transcripts, which represents 8.47 % of the total detected transcripts, 28045 are detected by all the three methods,16754 transcripts were detected by exactly two methods, shared between each method pair and a total of 23,061 transcripts were detected by a single method. (*Figure 3B*). The remaining majority of the detected transcripts (91.53%) do either not map the reference uninterruptedly or contain sequences that are marked as intronic, intergenic or unknown, to a varying degree. It is beyond the scope of this investigation to determine the validity of these results. The three sequencing methods and scmixology2 produced highly comparable isoform length distributions across all class codes (*Figure S3, Table S3*). The proportion of fully annotated transcripts (class code = “=“) is 13.3% in P29702_301, 16.7% in P29702_303, 13.1% in P29702_401, and 15.1% in scmixology2 (*Table S3*).

**Figure 3.**
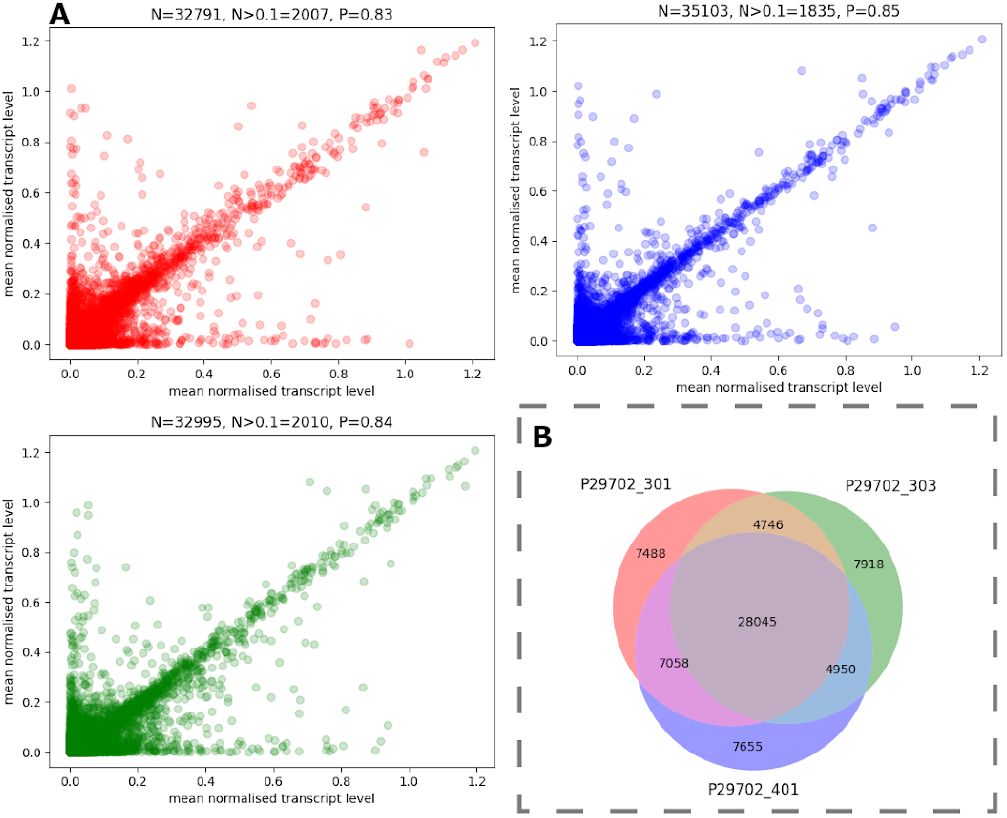
Isoform usage between all three libraries (PCS111 - P29702_301, FLT-seq - P29702_303 and PCA001 P29702_401). A) Comparative transcript expression levels between PCS111 vs. FLT-seq (red), FLT-seq vs. PCA001 (blue) and FLT-seq vs. PCA001 (green). B) Venn diagram for exact and complete intron-chain matching isoform presence in the three libraries, as defined by the “transcript” entry in the GENCODE annotation (human version 32).

Multi-transcript (isoforms) loci can give an indication of alternative splicing. In order of PCS111, FLT-seq, PCA001, we find the following number of multi-transcript loci: 13681, 14671, 12158,, which corresponds to roughly 1.3, 1.5, 1.4, transcripts per locus.

An example of a locus where several isoforms were identified is the XIST long noncoding RNA (*Figure 4*), where 5 different isoforms were detected by FLT-seq. XIST long noncoding RNA is indispensable for X-Chromosome inactivation and initiates the process early during development by spreading in cis across the X chromosome from which it is transcribed (Loda & Heard, 2019).

**Figure 4.**
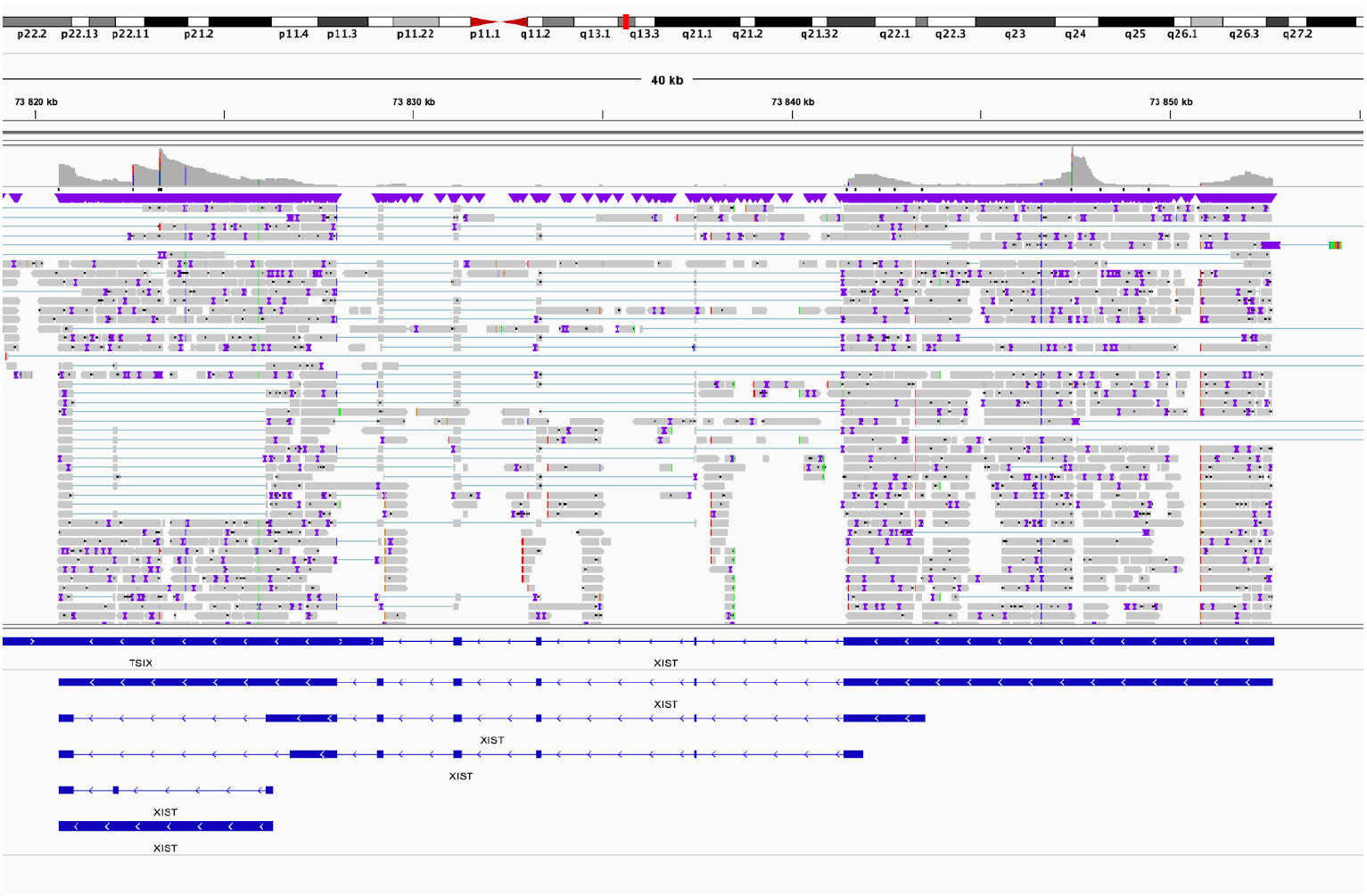
Isoform analysis of XIST long noncoding RNA. IGV view of FLT-seq data, Chromosome X: 73,820,656-73,852,723 (reverse strand) with 5 different transcripts detected (in order) ENST00000429829.6 ENST00000669898.1 ENST00000417942.5 ENST00000421322.3 ENST00000669898.1. In the middle of the graph, gray bars show aligned reads, gray histogram represents coverage. Lower part of the graph shows transcripts in blue: thin blue lines are intronic, solid blue lines stand for exonic regions.

### Full-length transcripts

In order to answer the question if any of the three methods are generating the full-length transcript data, we looked at gene body coverage (*Figure 5*). All three methods show a bias towards the 3’ end of the mRNA, which is likely due to a combination of a partly inefficient reverse transcription (which starts from the 3’ end) and the subsequent rounds of PCR amplification preferably amplifying short fragments (*Figure 5A*).

**Figure 5.**
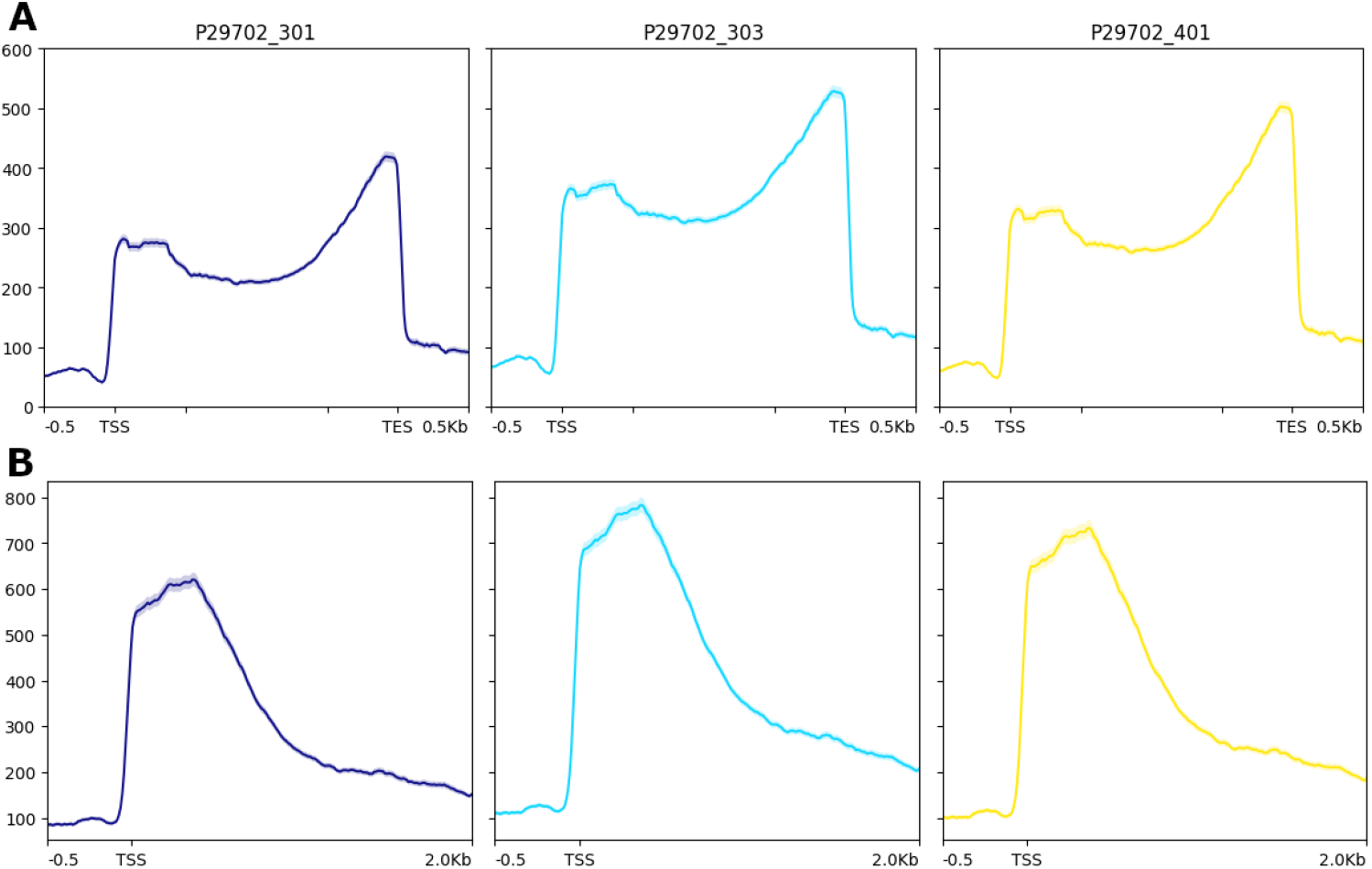
Gene coverage. Measured as mapping coverage to GENCODE 45 concatenated exons w. >0 coverage as measured by Deeptools2. Shown as A) in relative scale from TSS to TES and in B) in absolute scale from TSS to 2 Kb downstream. PCS111 - P29702_301, FLT-seq - P29702_303 and PCA001 P29702_401).

## Conclusion

The aim of this pilot study was to evaluate the performance of methods to add full-length mRNA isoform data to Chromium Single Cell experiments. The methods for sequencing full-length single-cell cDNA libraries were **FLT-seq** (Tian *et al* 2021), and two ONT protocols for Single-cell transcriptomics: cDNA-PCR sequencing kit **SQK-PCS111** or PCR expansion kit **EXP-PCA001 (***Figure 1C, Table S1*). Their performance was compared to publicly available data.

In terms of data, the three methods generated comparable results and all are suitable for isoform analysis. The identification of a substantial number of transcripts (28,045) that were consistently detected by all three methods reinforces the reliability of these findings (*Figure 3B*). This overlap builds confidence that these transcripts are likely real and biologically relevant, rather than artifacts of any single analysis pipeline. FLT-seq exhibited the highest percentage of fully annotated transcripts at 16.7% (*Figure S3, Table S3*). This may indicate a higher transcriptome coverage achieved through FLT-seq.

While the methods largely correlate in transcript levels, and many transcripts are detected by all three methods, there is no complete overlap. We are analyzing a limited sample number with low sequencing depth and do not have a reference dataset of “full-length” data to compare the methods. There are few available datasets to compare our data to.

We concluded that FLT-seq should be the method of choice due to the following differences:

1. FLT-seq and PCR expansion kit EXP-PCA001 use the latest R10 sequencing chemistry while cDNA-PCR sequencing kit SQK-PCS111 likely performed worse due to the obsolete R9 chemistry.
2. Compared to FLT-seq, PCR expansion kit EXP-PCA001 has a higher input requirement and requires more hands-on lab work with two rounds of PCR amplifications and bead purifications prior to library construction
3. FLT-seq is cost-effective to scale with off-the-shelf components.

In addition to overall isoform detection and barcode assignment performance, all tested methods exhibited a measurable proportion of reads with multiple TSO annotations, reflecting the presence of library preparation artifacts *(Table S2)*. The frequency of multi-TSO reads varied by protocol and serves as a key quality control metric for full-length single-cell cDNA sequencing. Minimizing such artifacts will be important for future method optimization and robust isoform discovery.

## Methods

### cDNA generation

GEM Generation & Barcoding was performed using the 10x Genomics Chromium Next GEM Single Cell 3’ Kit v3.1, following the manufacturer’s instructions (CG000315). 8300 K562 cells were loaded for GEM Generation on the Chromium X instrument (10x genomics) with a targeted cell recovery of 5000.

Following reverse transcription and bead cleanup, the barcoded cDNA library was amplified by PCR. For quality control and quantification, the bead-cleaned cDNA library was analyzed on the Agilent Bioanalyzer High Sensitivity Chip.

### Sequencing library generation and sequencing

The sequencing libraries for **FLT-seq, SQK-PCS111** and **EXP-PCA001** were generated from 5-10 ng of cDNA amplicons (*Table S1*.), following the original publication with modification (Tian *et al* 2021), see *supplementary method FLT-seq* or protocol from the manufacturer of the kit: *ONT_SQK-PCS111_single-cell-transcriptomics-10x-SST_v9148_v111_revE_12Jan2022-promet hion.pdf* and single-cell-transcriptomics-with-cdna-prepared-using-10x-SST_v9198_v114_revA_06Dec2023-p romethion (3).pdf and sequenced on the respective promethion flowcell (*Table S1*.)

## Supporting information

Supplementary Method FLT-seq

## Data analysis

The primary data analysis, i.e. resolving Chromium adapters and barcodes, read mapping and gene and transcript assignments, was performed using the “wf-single-cell” (v. 1.0.3) workflow from epi2me-labs (https://github.com/epi2me-labs/wf-single-cell). Further, the transcript expression matrices were normalized using the Scanpy (v. 1.10.1) python package and transcriptome annotations were manipulated using gffread (v. 0.12.7). The various scripts and commands that were run are found in a public git repository (see Data availability).

## Data availability

The sequence data has been deposited to ENA and can be found in accession PRJEB85328. Command-line scripts and jupyter notebooks containing source code for some of the figures in this publication can be found on GitHub along with supplementary figures (https://github.com/NationalGenomicsInfrastructure/NGI-ONT_scRNAseq-tech_note-2025)

## Acknowledgements

The authors would like to thank Rachel Thijssen for insights and discussions on FLT-seq, Gina Hendo for performing the GEM Generation & Barcoding, and Anniina Vihervaara for providing the K562 cells and allowing the usage of the single-cell cDNA libraries for this study.

The authors acknowledge support from the National Genomics Infrastructure in Stockholm funded by Science for Life Laboratory, the Knut and Alice Wallenberg Foundation and the Swedish Research Council, and SNIC/Uppsala Multidisciplinary Center for Advanced Computational Science for assistance with massively parallel sequencing and access to the UPPMAX computational infrastructure.

## Supplementary

Table S1 Methods for ONT library construction with enrichment of cDNA library.

Table S2 - multi TSO and multi R1

Table S3 - Classes of Transcripts for between all three libraries PCS111, FLT-seq, PCA001 and scmixology2 as classified by GFFCompare.

Figure S1 - Relative distance of TSO to read ends

Figure S2 - TSO hits in absolute distance from read ends

Figure S3 - Isoform length distribution comparison between PCS111, FLT-seq, PCA001 and scmixology2 and across different class codes.

Supplementary Method FLT-seq

