## Supplementary material for "Isoform-Level Analysis of 10x Genomics Single-Cell cDNA Libraries from Cultured K562 Cells Using Long-Read Sequencing": Table S1

**Table S1** - Methods for ONT library construction with enrichment of cDNA library.

| | **FLT-Seq**<br>single-cell full-length transcript sequencing by sampling | **SQK-PCS111**<br>Single-cell transcriptomics with cDNA prepared using 10x SST (2022) | **EXP-PCA001**<br>Single-cell transcriptomics with cDNA prepared using 10x SST (2023) |
|---|---|---|---|
| **Sample name** | **P29702_303** | **P29702_301** | **P29702_401** |
| Source | Tian *et al.* (2021) Genome Biology | Oxford Nanopore Technologies | Oxford Nanopore Technologies |
| Full-length cDNA enrichment | PCR amplification with specific primers | PCR amplification with specific primers & biotin pull-down | PCR amplification with specific primers & biotin pull-down |
| Primers | FPSfilA:<br>5’-ACTAAAGGCCATTACGGCCTACACGACGCTCTTCCGATCT-3’<br><br>RPSfilBr:<br>5’-TTACAGGCCGTAATGGCCAAGCAGTGGTATCAACGCAGAGTA-3’ | [Btn]Fwd_3580_partial_read1_defined<br>5'-/5Biosg/CAGCACTTGCCTGTCGCTCTATCTTCCTACACGACGCTCTTCCGATCT-3'<br>Rev_PR2_partial_TSO_defined<br>5'-CAGCTTTCTGTTGGTGCTGATATTGCAAGCAGTGGTATCAACGCAGAG-3'<br><br>cDNA Primer (cPRM)<br>Forward sequence:<br>5'-ATCGCCTACCGTGACAAGAAAGTTGTCGGTGTCTTTGTGACTTGCCTGTCGCTCTATCTTC-3’<br><br>cDNA Primer (cPRM)<br>Reverse sequence:<br>5’-ATCGCCTACCGTGACAAGAAAGTTGTCGGTGTCTTTGTGTTTCTGTTGGTGCTGATATTGC-3’ | [Btn]Fwd_3580_partial_read1_defined<br>5'-/5Biosg/CAGCACTTGCCTGTCGCTCTATCTTCCTACACGACGCTCTTCCGATCT-3'<br>Rev_PR2_partial_TSO_defined<br>5'-CAGCTTTCTGTTGGTGCTGATATTGCAAGCAGTGGTATCAACGCAGAG-3'<br><br>cDNA Primer (cPRM)<br>Forward sequence:<br>5'-ATCGCCTACCGTGACAAGAAAGTTGTCGGTGTCTTTGTGACTTGCCTGTCGCTCTATCTTC-3’<br><br>cDNA Primer (cPRM)<br>Reverse sequence:<br>5’-ATCGCCTACCGTGACAAGAAAGTTGTCGGTGTCTTTGTGTTTCTGTTGGTGCTGATATTGC-3’ |
| Input | 2.5-5 ng of cDNA amplicons | 10 ng of cDNA amplicons | 10 ng of cDNA amplicons |
| Based on ONT kit | Ligation Sequencing Kit | PCR-cDNA | Ligation Sequencing |

| | | | |
|---|---|---|---|
| | V14 (SQK-LSK114) | Sequencing Kit (SQK-PCS111) | Kit V14 (SQK-LSK114)<br><br>PCR Expansion (EXP-PCA001) for (cPRM) |
| Adapter | Ligation Adapter | Rapid Adapter T | Ligation Adapter |
| ONT flow cell | PromethION R10.4.1 | PromethION R9.4.1 | PromethION R10.4.1 |
