## Supplementary material for "Isoform-Level Analysis of 10x Genomics Single-Cell cDNA Libraries from Cultured K562 Cells Using Long-Read Sequencing": Table S2

**Table S2 - multi TSO and multi R1**

| **Multi-TSO search** | | | | |
|---|---|---|---|---|
| Number of TSOs per read | P29702_301 PCS111 | P29702_303 FLTseq | P29702_401 PCA001 | scmixology2 |
| 1 | 16096518 | 15801695 | 15789738 | 15851656 |
| 2 | 611120 | 2009971 | 347233 | 625671 |
| 3 | 29448 | 246154 | 13690 | 22379 |
| 4 | 2495 | 37547 | 3594 | 884 |
| 5 | 497 | 5737 | 1359 | 37 |
| 6 | 163 | 1336 | 718 | 4 |
| 7 | 86 | 394 | 461 | 0 |
| 8 | 50 | 214 | 304 | 0 |
| 9 | 26 | 145 | 190 | 1 |
| >=10 | 11 | 777 | 776 | 0 |
| **Multi-R1 adapter search** | | | | |
| Number of "read1-adapter" per read | P29702_301 PCS111 | P29702_303 FLTseq | P29702_401 PCA001 | scmixology2 |
| 1 | 15103792 | 13138542 | 15325604 | 13848633 |
| 2 | 488387 | 944466 | 315775 | 458027 |
| 3 | 18152 | 66568 | 7965 | 14824 |
| 4 | 790 | 5662 | 950 | 566 |
| 5 | 44 | 566 | 290 | 22 |
| 6 | 4 | 199 | 185 | 0 |
| 7 | 1 | 108 | 136 | 0 |
| 8 | 0 | 73 | 104 | 0 |
| 9 | 1 | 46 | 87 | 0 |
| >=10 | 0 | 186 | 547 | 0 |
