## Supplementary material for "Isoform-Level Analysis of 10x Genomics Single-Cell cDNA Libraries from Cultured K562 Cells Using Long-Read Sequencing": Table S3

**Table S3 - Classes of Transcripts for between all three libraries PCS111, FLT-seq, PCA001 and scmixology2 as classified by GFFCompare.**

| | **P29702_301 PCS111** | | **P29702_303 FLTseq** | | **P29702_401 PCA001** | | **scmixology2** | |
|---|---|---|---|---|---|---|---|---|
| Transcript Classes | count | % | count | % | count | % | count | % |
| = | 28995 | 13.3 | 30247 | 16.7 | 28122 | 13.1 | 29507 | 15.1 |
| c | 5002 | 2.3 | 3152 | 1.7 | 4895 | 2.3 | 2893 | 1.5 |
| e | 5670 | 2.6 | 3909 | 2.2 | 5722 | 2.7 | 5195 | 2.7 |
| i | 87243 | 40.0 | 53144 | 29.3 | 81870 | 38.2 | 77061 | 39.5 |
| j | 17307 | 7.9 | 16392 | 9.0 | 18073 | 8.4 | 21240 | 10.9 |
| k | 4608 | 2.1 | 6536 | 3.6 | 5061 | 2.4 | 6760 | 3.5 |
| m | 3231 | 1.5 | 5207 | 2.9 | 3534 | 1.6 | 2352 | 1.2 |
| n | 5786 | 2.7 | 7772 | 4.3 | 6191 | 2.9 | 4408 | 2.3 |
| o | 3918 | 1.8 | 3845 | 2.1 | 4373 | 2.0 | 3550 | 1.8 |
| p | 3355 | 1.5 | 2527 | 1.4 | 3211 | 1.5 | 2413 | 1.2 |
| s | 1684 | 0.8 | 5694 | 3.1 | 2205 | 1.0 | 770 | 0.4 |
| u | 40150 | 18.4 | 34403 | 19.0 | 40320 | 18.8 | 29909 | 15.3 |
| x | 10882 | 5.0 | 8154 | 4.5 | 10660 | 5.0 | 9056 | 4.6 |
| y | 275 | 0.1 | 207 | 0.1 | 243 | 0.1 | 165 | 0.1 |
| Total | 218106 | | 181189 | | 214480 | | 195279 | |
