## Supplementary material for "Isoform-Level Analysis of 10x Genomics Single-Cell cDNA Libraries from Cultured K562 Cells Using Long-Read Sequencing": Figure S1

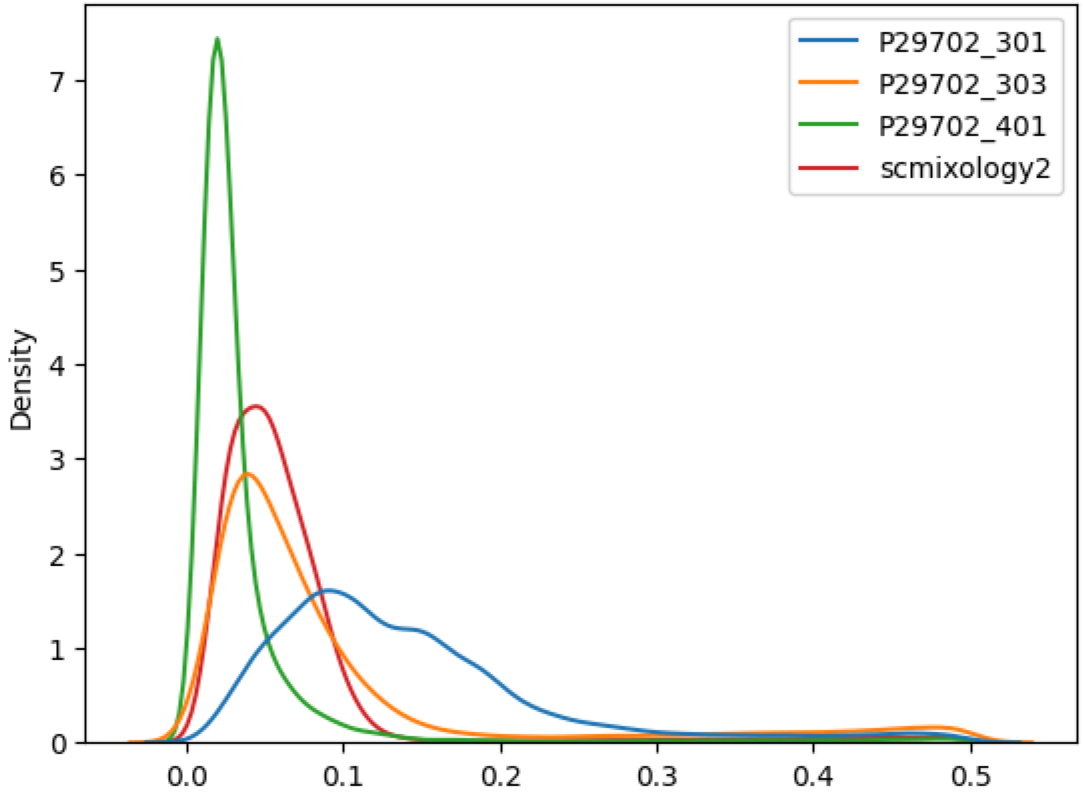


**Figure S1** - Relative distance of TSO to read ends, i.e. 0.0 is 3’ or 5’ end and 0.5 is the middle, in cDNA libraries shown as kernel density estimates. Green: PCA001, Orange: FLT-seq, Blue: PCS111. PCA: TSO mostly at end
