## Supplementary figures and images for "Isoform-Level Analysis of 10x Genomics Single-Cell cDNA Libraries from Cultured K562 Cells Using Long-Read Sequencing"

### Figure S2

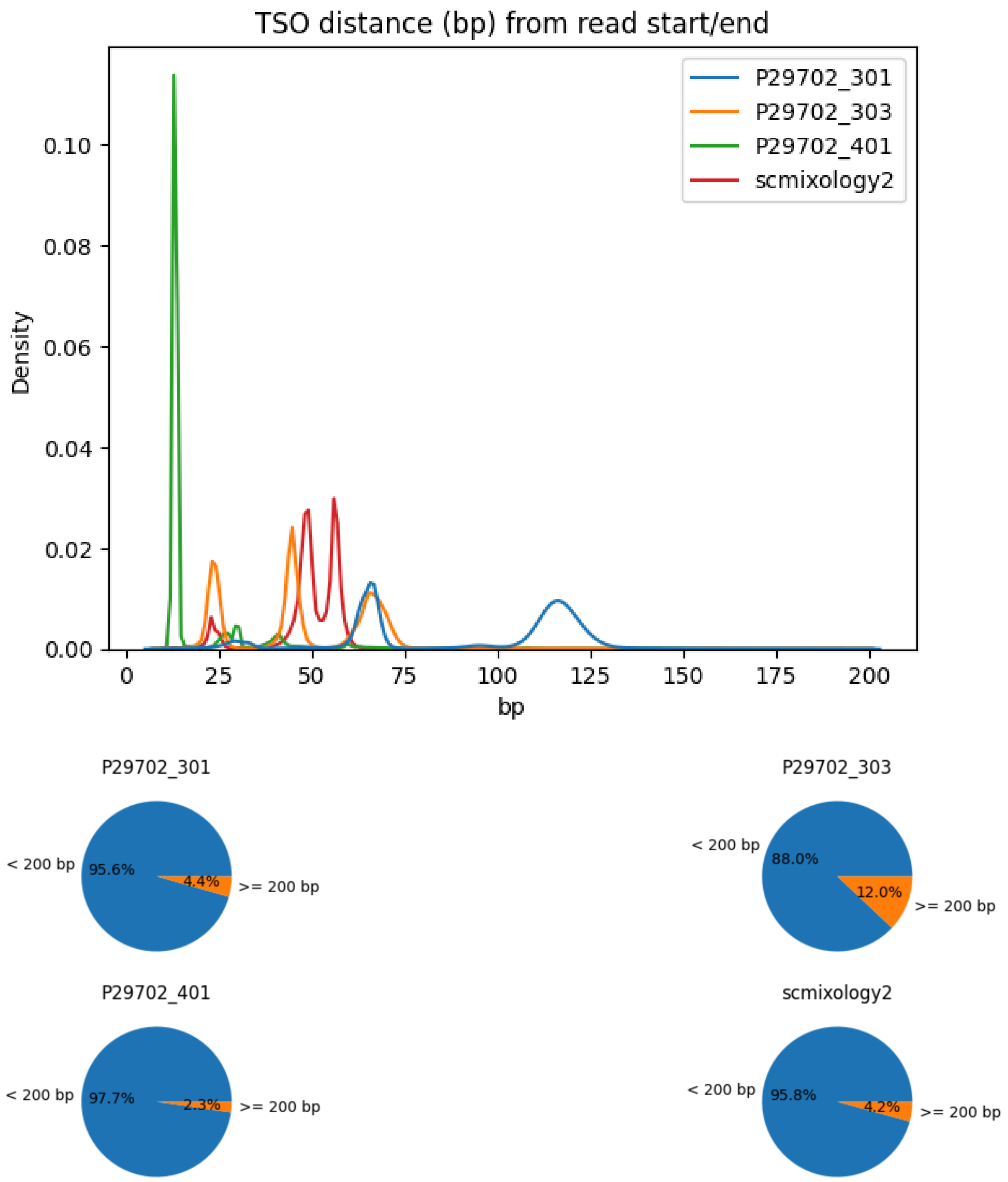


**Figure S21 - TSO hits in absolute distance from read ends**
