## Supplementary material for "Isoform-Level Analysis of 10x Genomics Single-Cell cDNA Libraries from Cultured K562 Cells Using Long-Read Sequencing": Figure S3

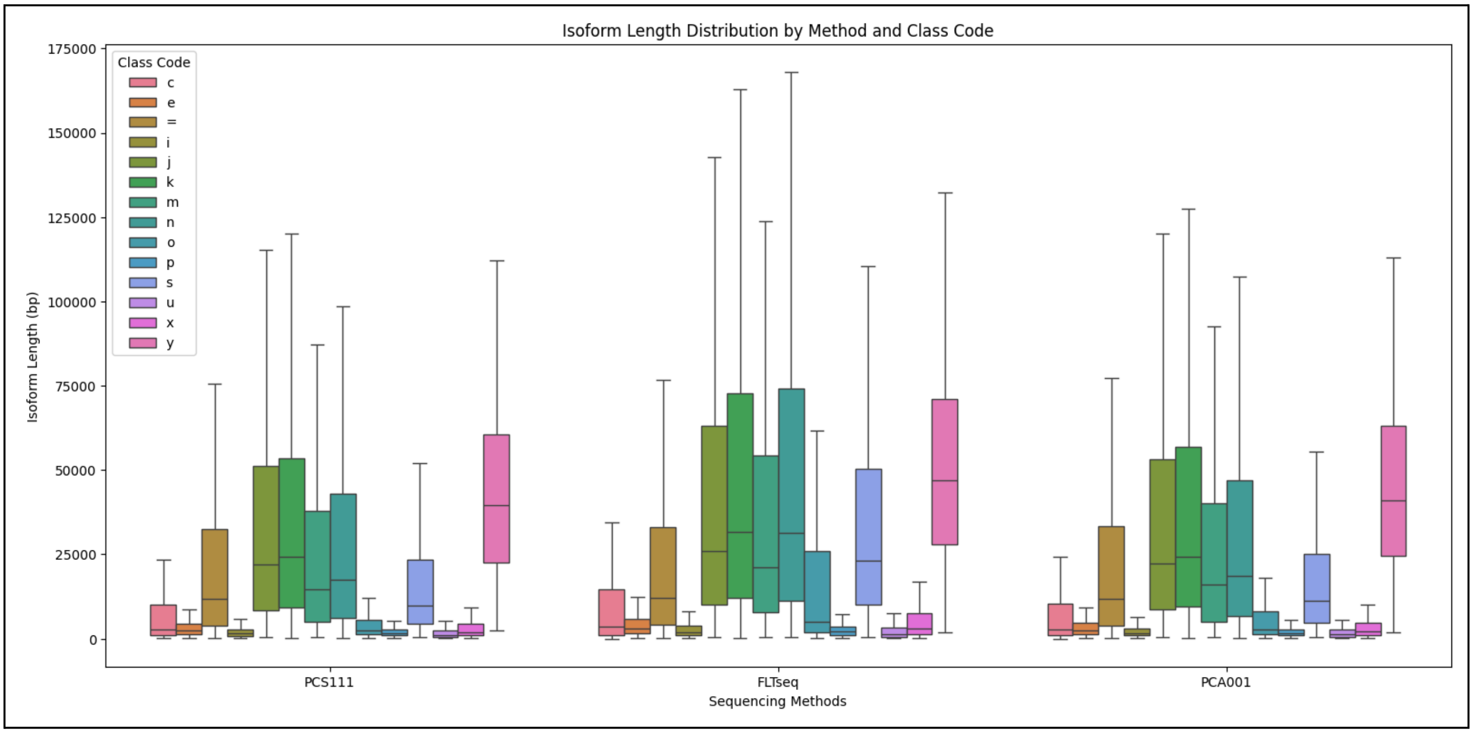


**Figure S3 - Isoform length distribution comparison between PCS111, FLT-seq, PCA001 and scmixology2 and across different class codes.**

**Figure Description -**

| Code | Description |
|---|---|
| = | Complete, exact match of intron chain |
| c | Contained in reference (intron compatible) |
| k | Containment of reference (reverse containment) |
| m | Retained intron(s), all introns matched or retained |
| n | Retained intron(s), not all introns matched/covered |
| j | Multi-exon with at least one junction match |
| e | Single exon transfrag partially covering an intron, possible pre-mRNA fragment |
| o | Other same strand overlap with reference exons |
| s | Intron match on the opposite strand (likely a mapping error) |
| x | Exonic overlap on the opposite strand (like o or e, but on the opposite strand) |
| i | Fully contained within a reference intron |

| | |
|---|---|
| y | Contains a reference within its intron(s) |
| p | Possible polymerase run-on (no actual overlap) |
| r | Repeat (at least 50% bases soft-masked) |
| u | None of the above (unknown, intergenic) |
