## Supplementary Method FLT-seq for "Isoform-Level Analysis of 10x Genomics Single-Cell cDNA Libraries from Cultured K562 Cells Using Long-Read Sequencing"

The cDNA from the Chromium Next GEM Single Cell 3' Reagent Kit v3.1 was used to generate the Oxford Nanopore Technology (ONT) long-read sequencing libraries, according to a modified FLT-seq protocol (ref bellow) and the SQK-LSK114 Ligation Sequencing Kit (Oxford Nanopore Technologies, Oxford, UK).

1) PCR amplification. For each sample, 5 PCR-reactions in parallel were set up with the following reagents: 10 µl 5x PrimeSTAR GXL Buffer (Takara Bio Inc., Kasatsu, Japan), 4 µl 2.5 mM dNTP solution (New England Biolabs, Ipswich, USA), 1 µl 10 µM FPSfilA Primer (FPSfilA: 5'-ACTAAAGGCCATTACGGCCTACACGACGCTCTTCCGATCT-3', Thermo Fisher Scientific, Waltham, USA), 1 µl 10 µM RPSfilBr Primer (RPSfilBr: 5'-TTACAGGCCGTAATGGCCAAGCAGTGGTATCAACGCAGAGTA-3', Thermo Fisher Scientific, Waltham, USA), 1 µl PrimeSTAR GXL Polymerase (Takara Bio Inc., Kasatsu, Japan).

Samples were PCR-amplified with the following program: 98°C for 30 sec, 8 cycles of (98°C for 10 sec, 65°C for 15 sec, 68°C for 8 min), 68°C for 10 min, 10°C hold.

2) Bead clean up. 5 x 50 µl of PCR product were pooled for each sample, mixed with 200 µl (0.8x) AMPure XP beads (Beckman Coulter Inc., Brea, USA) and incubated at room temperature for 10 minutes. The mix was placed on a magnet and the clear supernatant was removed. With the tubes still on the magnet, 200 µl 80% ethanol was added. The ethanol was removed and the wash with 200 µl 80% ethanol was repeated once for a total of two washes. Beads were air dried for 1 minute. The beads were resuspended in 51 µl Buffer EB (Qiagen N.V. Hilden, Germany) off the magnet and incubated at room temperature for 5 minutes. Placed back on the magnet, 50 µl of the clear supernatant was removed and transferred to a new tube.

3) Quality control. The amplified cDNA was quantified using Qubit dsDNA HS assay (Thermo Fisher Scientific, Waltham, USA) and Fragment Analyzer HS NGS Fragment Kit (Agilent Technologies Inc., Santa Clara, USA).

4) Long-read sequency library preparation. 200 ng of the amplified cDNA from the previous step was used as an input for library preparation according to SQK-LSK114 Ligation Sequencing Kit (Oxford Nanopore Technologies, Oxford, UK) with following modifications (original values ~~crossed out~~): *3. DNA repair and end-prep*: 3.6 Using a thermal cycler, incubate at 20°C for ~~5~~ 20 minutes and 65°C for ~~5~~ 20 minutes. 3.10 Incubate on a Hula mixer (rotator mixer) for ~~5~~ 15 minutes at room temperature. *4. Adapter ligation and clean-up* 4.9 Add ~~40~~ 50 µl of resuspended AMPure XP Beads (AXP) to the reaction and mix by flicking the tube.

5) Long-read sequencing. We sequenced 50 fmol of library per sample in a Promethion R10.4.1 flowcell, according to SQK-LSK114 Ligation Sequencing Kit (Oxford Nanopore Technologies, Oxford, UK).

ref FLT-seq:

Tian, L., Jabbari, J.S., Thijssen, R. et al. (2021). Comprehensive characterization of single-cell full-length isoforms in human and mouse with long-read sequencing. Genome Biol 22, 310 . https://doi.org/10.1186/s13059-021-02525-6
